# Tension-area relationship in compartmentalized crumpled plasma membrane: a mechanistic model and its implications

**DOI:** 10.1101/2025.01.04.628534

**Authors:** Andrey K. Tsaturyan

## Abstract

The plasma membrane is a liquid lipid bilayer containing both dissolved proteins and proteins anchoring the membrane to the underlying actin cortex. Membrane tension, a 2D analog of pressure in a 3D liquid, is believed to play a crucial role in organizing essential processes within cells and tissues. This, along with recent, conflicting data on the speed of membrane tension propagation, highlights the need for a comprehensive mechanical model to describe tension in the cortex-anchored plasma membrane as a function of transmembrane hydrostatic pressure difference and excess membrane area due to cortex contraction. In this study, we present a mechanical model of plasma membrane compartments, separated by “picket fences” of cortex-anchoring proteins permeable to lipids. Beyond hydrostatic pressure, the model incorporates the 2D osmotic pressure exerted by membrane-dissolved proteins. Our findings reveal that the tension-area relationship within a membrane compartment exhibits a seemingly paradoxical feature: in a specific range of membrane surface area, an increase in area leads to a rise in tension. We further model the tension-area relationship for an ensemble of membrane compartments, which exchange membrane area through shared borders, and discuss potential biological implications of this model.

## Introduction

The plasma membrane serves as a barrier, separating the cell’s interior from its external environment. It primarily consists of a lipid bilayer and lipid-associated proteins. Some of these proteins are dissolved in the lipids and can diffuse along the membrane surface, while others anchor the membrane to the underlying cortex (Singer, Nicolson 1972; Mayor et al. 2023). The cortex is composed of actin filaments, actin-binding proteins that form the actin meshwork, and the motor protein myosin, which enables its ATP-dependent active contraction (Chugh, Paluch 2018). This contraction tends to crumple the membrane through its anchoring proteins. Changes in the cortex, such as optogenetically induced rapid growth of cell protrusions or localized cortex contraction, lead to quick changes in membrane tension estimated by measuring the force required to hold membrane tethers pulled from different locations on the cell surface (De Belly et al. 2023).

Mechanically, the membrane behaves like a 2D viscous liquid, allowing lipids to flow along its surface. Membrane tension in the lipid bilayer acts as the 2D equivalent of pressure in a 3D liquid (but with the opposite sign), representing the force per unit length along the membrane’s edge – positive when the force pulls the membrane and negative when it pushes. Additionally, since the cortex is likely permeable to cytoplasm, the membrane is subjected to a pressure difference between the intra- and extracellular spaces. The membrane also resists changes in its curvature through bending torque (Evans, Skalak 1980; Kozlov 2006; Zucker et al. 2023).

The tension propagates very quickly between two tethers pulled from a membrane bleb (Shi et al. 2018; De Belly et al. 2023). A bleb is a bubble-like protrusion of the cell membrane caused by intracellular pressure, typically resulting from partial detachment of the cell membrane from the underlying cortex, cortex rupture, or active contraction (Charras 2008). In contrast, membrane tension propagation between two tethers pulled from cortex-attached membrane of the same cells is much slower, if it occurs at all (Shi et al. 2018; De Belly et al. 2023). The tension propagation velocity varies depending on the cell type and even the surface region from which the tether is pulled. For example, tension propagates quickly along the axon but slowly along dendrites in neurons (Shi et al. 2022). Models developed to address the problem and explain the contradictorily data are based on the assumption that the plasma membrane has a stretching modulus of 40-100 pN/μm (Shi et al. 2018; Cohen, Shi 2020; De Belly et al. 2023). This estimate implies that a moderate membrane tension of 0.04-0.1 mN/m would cause a two-fold increase in the actual membrane surface area or the area per lipid molecule which seems unrealistic. Indeed, these estimates are at least three orders of magnitudes higher than the widely accepted value of 10^5^ pN/μm (Evans, Skalak 1980). This more realistic estimate suggests that even a very high membrane tension of 0.5 mN/m, which leads to membrane rapture (Tan et al. 2011), results in only a 0.5% increase in the actual membrane area.

These controversial data, along the crucial role that membrane tension plays in the spatial coordination of various essential physiological processes within cells and tissues (Diz-Muñoz et al. 2013; Sens, Plastino 2015; Pontes et al. 2017; Sitarska, Diz-Muñoz 2020; Ghisleni, Gauthier 2024) highlight the need for a comprehensive mechanical model of plasma membrane tension. Such a model must account for the membrane’s anchoring in the cortex, its crumpling due to cortical contraction, and the hydrostatic pressure difference.

A mechanistic model of a plasma membrane compartment, surrounded by picket fences formed by cortex-anchoring proteins, was suggested (Barnoy et al. 2023). The compartmentalized membrane model, according to which the plasma membrane is divided into distinct compartments via these picket fences, has been validated by multiple studies (Fujiwara et al. 2002, 2016; Kuzumi et al. 2005, 2011, 2012; Jacobson et al, 2019; Mayor et al. 2023). The average size of these compartments is estimated at 200-230 nm, with a standard deviation of *ca*. 50-70 nm (Kuzumi et al. 2012). In our study (Barnoy et al. 2023), we accounted for lipid flow between neighboring compartments across their borders, leading to membrane area redistribution along the cell surface and membrane tension equilibration in neighboring compartments. The mechanical properties of the membrane in this model were simplified: it was considered inextensible, with bending stiffness neglected, so that tension was entirely determined by hydrostatic pressure and surface curvature according to Laplace’s law. An advanced model (Dharan et al. 2025) incorporates membrane bending stiffness, though it was constrained to cases with small membrane excess area to allow for an analytical solution of the model’s linear equations. In that model, the relationship between membrane tension and excess membrane area at a constant transmembrane pressure difference is described analytically using Bessel functions. The relationship is monotonic: as excess membrane area approaches zero, membrane tension approaches infinity, while at higher membrane excess area, tension asymptotically approaches a constant level determined by the boundary conditions at the compartment border. The model also accounts for the redistribution of the membrane area between the compartments driven by tension-induced membrane flow across compartment borders. It further predicts that tension propagation follows a diffusion-like equation, where the diffusion coefficient is proportional to the pressure difference, *P*, permeability of the compartment borders, *λ*, square of the compartment size, *a*, and the normalized membrane excess area, *β*, in power 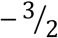. The model’s prediction that an increase in the pressure difference would accelerate tension propagation was confirmed experimentally (Dharan et al., 2025). The model offers a plausible explanation for the wide range of tension propagation velocities observed experimentally.

In this work, we present a more general non-linear model for a membrane compartment and an ensemble of membrane compartments capable to exchanging membrane surface area across their borders. We modeled an axisymmetric membrane compartment and numerically calculated the relationship between membrane tension at its circular border and the membrane’s excess area, under a constant pressure difference and various boundary conditions. Additionally, we considered the potential contribution of the 2D osmotic pressure from membrane-dissolved proteins on the tension-area relation. Our results reveal a seemingly paradoxical tension-area diagram for a membrane compartment: within a certain range of membrane excess area, the compartment exhibits “negative elasticity” where increasing the membrane area leads to rise in tension. Furthermore, we show that an ensemble of membrane compartments with this behavior acts as a membrane reservoir, maintaining nearly constant tension over a range of variations in excess membrane area.

## Model description

### Membrane mechanics

We consider membrane as a 2D surface with the total curvature (the trace of the curvature tensor or the sum of the two principal curvatures), *J*, and the Gaussian curvature (the determinant of the curvature tensor of the product of the principal curvatures), *K*. The static equilibrium equations for the normal and tangential components of membrane stress are as follows (Zucker et al. 2023)

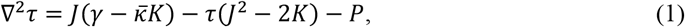

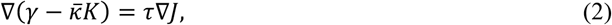

respectively. Here *γ* is membrane tension, 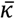 is a constant Gaussian bending modulus, τ is membrane torque, ∇ is the 2D covariant gradient operator, and ∇^2^ is the 2D Laplace operator along the membrane surface. Neglecting spontaneous membrane curvature, a linear elastic constitutive equation for the bending torque can be written as (Kozlov 2006; Zucker et al. 2023):

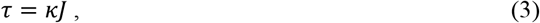

Where κ is a constant membrane bending stiffness.

Integration of Eq. (2) results in

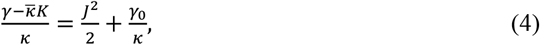

where *γ*_0_ is an integration constant that is membrane tension in a flat reservoir that is in equilibrium with the system.

### Axisymmetric compartment

For simplicity, assume axial symmetry of the membrane compartment. Then all variables depend only on the natural parameter, *s*, the distance along the line of intersection of a plane passing through the symmetry axis and the membrane surface. Then the total, *J*, and Gaussian, *K*, curvatures in Eq. (1), and the 2D Laplace operator are

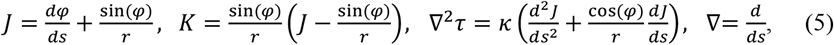

where *φ* is the angle between the tangent to the curve and a horizontal plane, and *r* is radial coordinate in a cylindrical coordinate system (*r, θ, z*).

The system of the equilibrium equations, Eqs (1-3), becomes

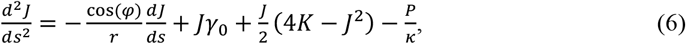

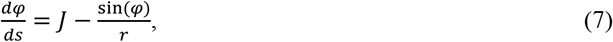

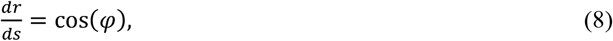

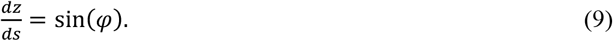

### Boundary conditions

The boundary conditions at the dome center are

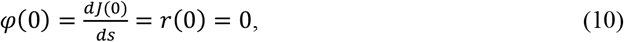

which are the conditions of axial symmetry and zero sheer force at the compartment pole. At the dome boarder, *r* = *a*, we use two extreme boundary conditions:

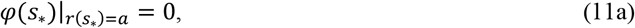

called here ‘stiff’ or zero angle conditions, or

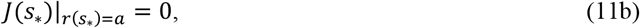

“soft’ or zero torque conditions. The condition Eq. (11a) provides the most constraint condition – the membrane tangent is parallel to surface of underlying cortex. The zero torque condition Eq. (11b) is the most “soft” and less constrained. The precise mechanical properties of the interface between lipid membrane and cortex-linking proteins are unknown. Nevertheless any physically plausible boundary conditions are within a range between two extremes (11a) and (11b).

### Dimensionless variables and equations

Introduce dimensionless variables and parameters:

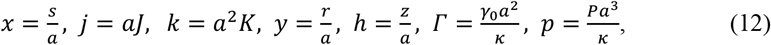

and rewrite the system of ODE, Eqs. (6-9), in dimensionless form:

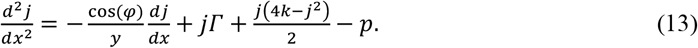

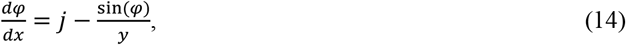

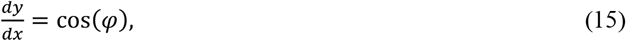

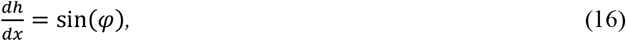

where 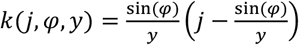 with boundary conditions

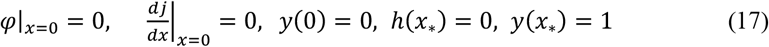

at the dome center and

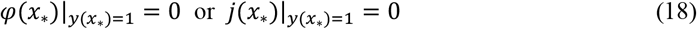

at the compartment border. The first and second formulas in Eq. (18) correspond to the stiff and soft conditions defined by Eqs. (11a-11b), respectively.

### Numerical implementation

The system of ordinary differential equations (ODE), Eqs. (13-16), was solved numerically at a given dimensionless pressure difference *p* and tension *Γ* with boundary conditions Eq. (17) at the pole, a condition at the border Eq. (18), plus conditions for *y, h*: *y*(0) = 0, *y*(*x*_∗_) = 1, *h*(*x*_∗_) = 0 using a 4th order implicit Runge-Kutta method. The “shooting” method was used to satisfy the boundary conditions. An initial value of *j*(0) was chosen and *j, φ, y, h* were calculates as functions of *x* until *y*(*x*) reached 1. Then curvature at the pole, *j*(0), was adjusted and integrations were repeated until the ‘stiff’ or ‘soft’ boundary condition (18) was satisfied at a given precision of 10^−3^.

Alternatively, the dome shape was calculated for a given normalized hydrostatic pressure, *p*, and the normalized membrane excess surface area, β, in power 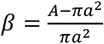,using the energy minimization software, Surface Evolver, based on the finite element method (Brakke 1992). Here *A* represents the membrane’s surface area within the compartment, and *πρ*^2^ is the membrane’s projection area. In this case, dimensionless membrane tension *Γ* was calculated as the Lagrange multiplier for constraint on the normalized membrane surface area. At least 300 finite elements were used; the energy minimizing was repeated until the relative change in energy and *Γ* per an iteration was ≤10^−3^. The results obtained by these two numerical methods for moderate *β* values were the same with a relative precision of 10^−3^.

## Results

The calculated membrane shapes, obtained at a constant normalized pressure *p* = 20 and under two different boundary conditions at the border (Eq. 18), are illustrated in Fig. 1.

**Fig. 1.**
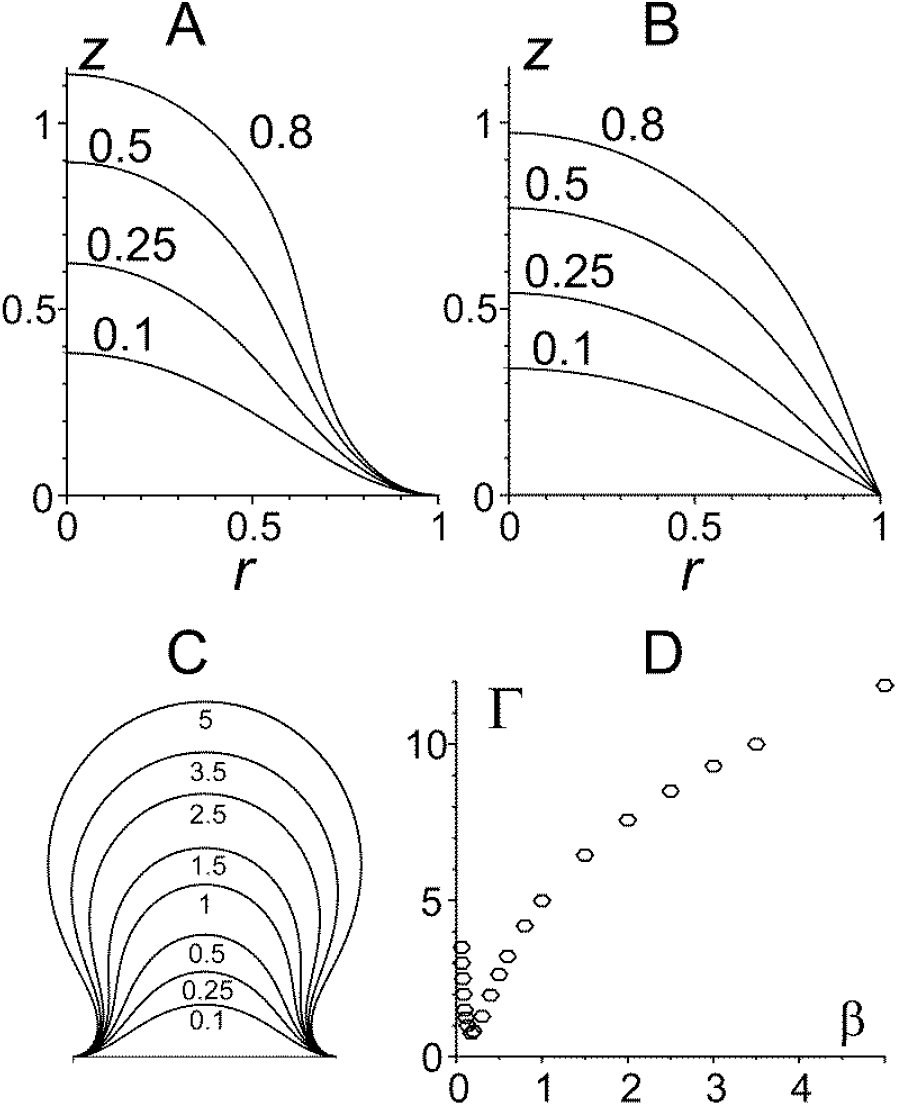
*A, B*: calculated membrane shapes at a constant normalized pressure *p* = 20 with ‘stiff’ (*A*) and ‘soft’ (*B*) boundary conditions at various normalized excess membrane surface areas (shown next to the plots); *C*: same as A at a wider range of normalized membrane excess area; *D*: normalized membrane tension, Γ, as a function of normalized membrane excess area, *β*, for the data set *C*.

As expected, at the same normalized membrane excess area, membrane domes under ‘stiff’ conditions was slightly higher than those under ‘soft’ conditions, while maintaining the same membrane excess area (Fig. 1*A, B*).

At elevated normalized membrane excess areas, *β*, the tip and central part of the dome exhibit a nearly spherical shape with a transition region next to the compartment border (Fig. 1*C*). The tension-area diagram (Fig. 1*D*) displays a non-monotonic and U-shaped pattern: tension decreases with *β* until it falls below a critical value, *β*_*c*_ and then increases when *β* exceeds *β*_*c*_. The normalized tension, *Γ*_*c*_, corresponding to the critical membrane excess area *β*_*c*_ represents the minimal membrane tension at a given normalized pressure, *p*.

Fig. 2 depicts the numerically calculated normalized membrane tension Γ across various normalized membrane excess areas, *β*, and normalized pressure differences, *p*, under both ‘stiff’ and ‘soft’ boundary conditions.

**Fig. 2.**
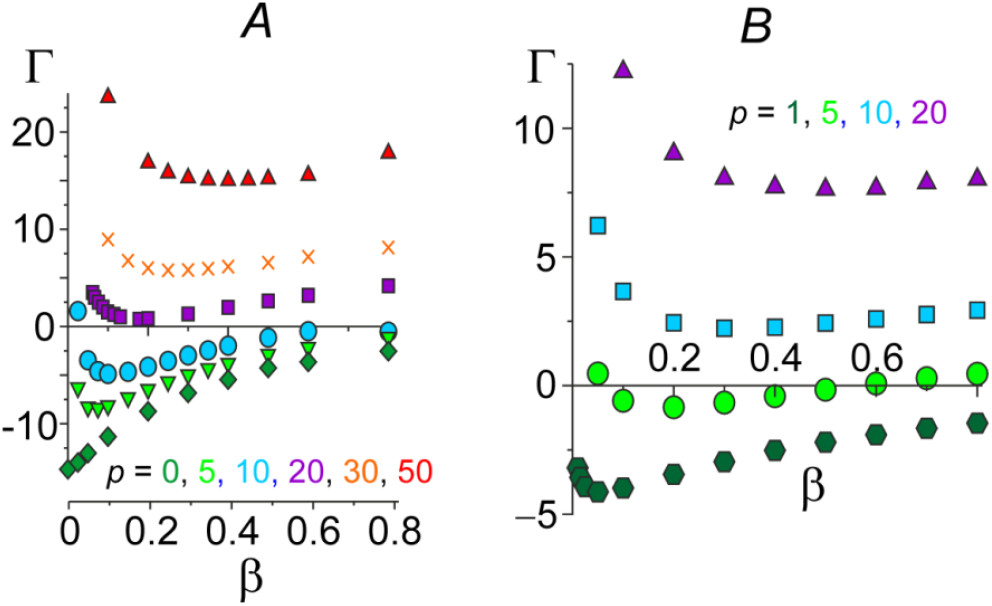
The dependence of normalized membrane tension, *Γ*, on normalized membrane excess area, *β*, in an axisymmetric membrane dome at different intracellular normalized pressures, *p* (denoted by different symbols and colors) and ‘stiff’ (*A*) and ‘soft’ (*B*) boundary conditions.

The tension-area diagrams exhibit a non-monotonic trend for all pressures tested under both types of boundary conditions. The critical membrane excess area, *β*_*c*_, and the critical (minimal) normalized tension, *Γ*_*c*_, increase with an elevation in the normalized transmembrane pressure *p* (Fig. 2*A, B*). Notably, under soft boundary conditions, both the normalized critical membrane surface area *β*_*c*_ and tension *Γ*_*c*_ were higher compared to stiff conditions (Fig. 2*A, B*).

In scenarios where there is no pressure difference, a threshold negative compressive tension is necessary to induce membrane buckling and push an excess membrane area into the compartment. Until this threshold is surpassed, the membrane remains flat, signifying *β*=0. For stiff boundary conditions, the absolute value of this negative tension threshold is higher than that for soft conditions (Fig. 2). Analytically derived from a linearized model, the threshold tension values are approximately –5.78 for soft and –14.68 for stiff conditions (Dharan et al. 2025).

The tension-area diagram at zero transmembrane hydrostatic pressure, *p* = 0 (green diamonds in Fig. 2*A*), can be considered as a limiting case of the non-monotonic diagrams at higher pressure difference with, featuring a degenerate vertical descending branch. As *p* increases from zero to infinity, the critical normalized excess area, *β*_*c*_, ascends from zero to 1. The very high normalized pressure difference at a moderate absolute pressure difference *P* corresponds to negligibly small membrane bending stiffness or torque-less membrane, as per Eq. (12).

### Physical nature of non-monotonic tension-area diagram

To gain a qualitative understanding of the intriguing non-monotonic dependence of Γ on *β*, we consider a simplified model of a torque-less membrane with vanishing bending stiffness. In this scenario, the membrane surface takes on a spherical shape, and pressure and tension adhere to the Laplace law:

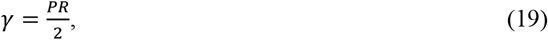

where *R* is the membrane radius. According to Eq. (19), membrane tension at a constant pressure is solely determined by the radius of the membrane sphere. Eq. (19) implies that tension in two spherical membrane segments with identical radii and differing height and surface area will be identical. Fig. 3*A* illustrates two spherical membrane domes with the base radius, *a*, identical radii, *R*, and pressure *P* but varying surface areas. As per from Eq. (19), the tension in these domes should be identical. The resulting tension-area diagram at a constant pressure is depicted in Fig. 3*B*. This non-monotonic tension-area diagram serves as an explanatory model for well-known behavior of a soap bubble when blown out of a round hole – as the bubble approaches the hemisphere, it breaks off and takes away.

**Fig. 3.**
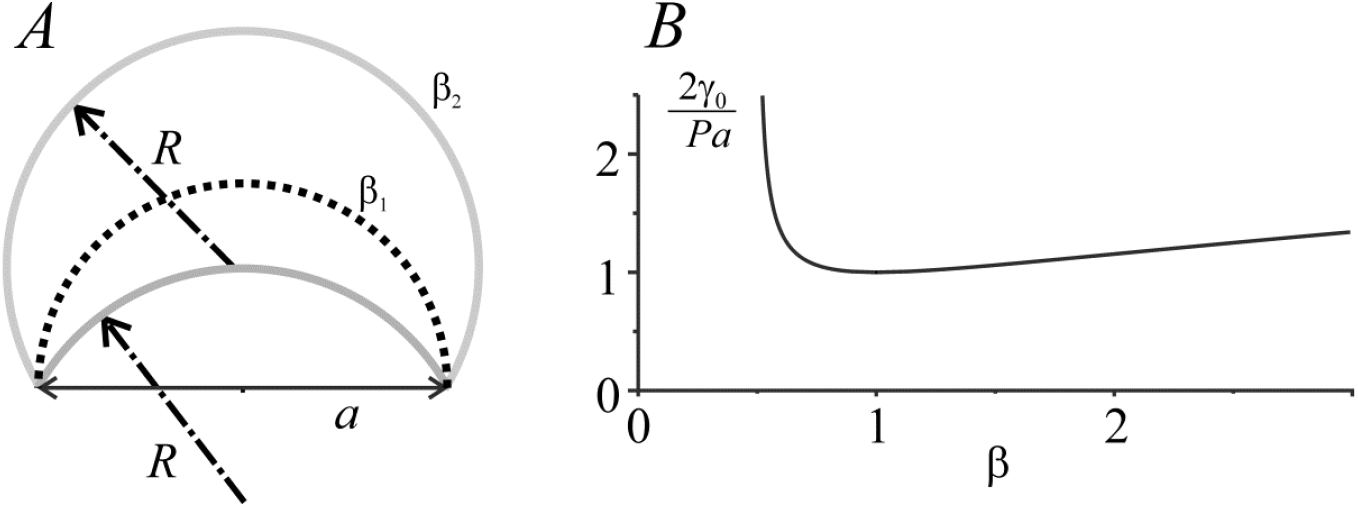
*A*: schematic representation of simplified torque-less membrane spherical domes under the same intracellular pressure and base radius, *a*; *β*_1_ < 1, *β*_2_>1 denote normalized membrane excess areas of two spherical ‘domes’ with the same radii *R* and, consequently, the same tension. A hemisphere with *R* = *a* and *β* = 1 is indicated by a dotted line. *B*: the dependence of normalized tension, *Γ*, on *β* at a normalized pressure difference of *p* = 2 for the torque-less membrane.

In Fig. 3, the critical membrane dome radius is equal to the base radius, *a*, and the tension-area diagram *γ*(*β*) exhibits a minimum at *β* = *β*_*c*_ = 1 with the surface area *A*_*m*_ = 2π*a*^2^ and the base area is *A*_*b*_ = π*a*^2^. For domes with *β*_1_ < 1 or *β*_2_>1 the spherical radii are higher resulting in higher membrane tension, *γ*.

The introduction of membrane bending stiffness does not qualitatively alter the tension-area diagram, although it reduces the critical excess surface area *β*_c_ (Fig. 2). The shift in *β*_c_ is more pronounced for stiff boundary condition (Fig. 2*A*) than for soft one (Fig. 2*B*). The lower the intracellular pressure, the lower is *β*_c_ (Fig. 2). As the pressure difference approaches zero, *β*_c_ likewise approaches zero (Fig. 2). At a non-zero pressure difference the non-monotonic tension-area diagram is between tow limit cases: i) the pressure-driven soap film-like mechanism, as described by Eq. (19) and illustrated by Fig. 3*B*; ii) the non-linear loss of stability of a flat elastic membrane loaded by compressing tension at (*p* = 0, Fig. 2*A*).

### Steady-state tension in an ensemble of membrane compartments

To gauge the potential repercussions of the peculiar tension-area diagram, consider first two adjacent membrane compartments under a constant intracellular pressure. The membrane flow rate, *q*, per a unit length the border between the compartments is driven by a tension difference between the compartments, Δ*γ* = *γ*_1_ − *γ*_2_, where *γ*_1_, *γ*_2_ are membrane tensions at the border in the compartments 1 and 2. A theory of flow of 2D viscous membrane through a regular picket fence of membrane anchored proteins (Barnoy et al. 2023; Dharan et al., 2025) gives

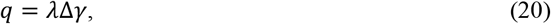

where λ is the border permeability proportional to the membrane viscosity. The permeability λ depends on the ratio of a picket diameter and the distance between neighbor pickets. The membrane flow is directed to the compartment with higher tension.

Assume the two compartments possess equal base areas and share a permeabilized border, enabling membrane flow between them. For simplicity, assume the total normalized membrane area in both compartments remains constant. If the sum of the excess membrane areas in the two compartments *β*_1_+*β*_2_ < 2*β*_*c*_ (*β*_1_ and *β*_2_ represent the excess membrane areas in the first and second compartment, respectively), the state where the areas are constant and equal to each other is stable because a perturbation of this state results in an increase in tension in the compartment with lower membrane area and subsequent membrane sucking to this compartment from its neighbor (Fig. 4*A*). If *β*_1_+*β*_2_>2*β*_*c*_, the state with equal membrane areas becomes unstable. A minor difference in initial tensions would lead to the influx of membrane into the compartment with higher tension. The steady equilibrium in the membrane redistribution corresponds to unequal membrane areas at equal tensions: one compartment stabilizes on the left descending branch of the diagram, while another, with a higher membrane area stabilizes on the right ascending branch of the diagram (Fig. 4*B*).

**Fig. 4.**
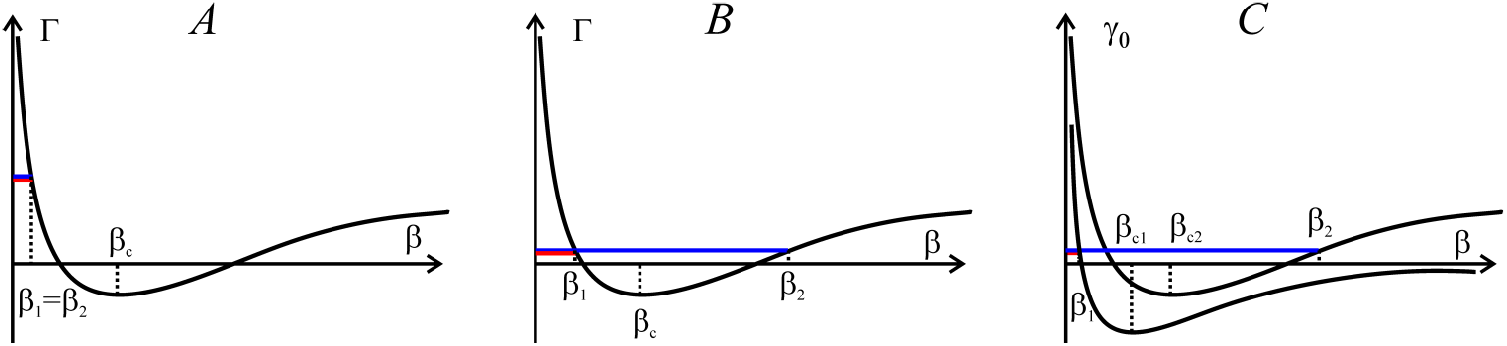
Illustration of membrane area distribution between two compartments. *A*: two compartments of the same base radii with the total excess membrane areas, *β*_1_+*β*_2_ < 2*β*_*c*_. Stable equilibrium is achieved at equal membrane areas, *β*_1_ = *β*_2_ < *β*_*c*_. *B*: similar to *A* with *β*_1_+*β*_2_>2*β*_*c*_. Stable equilibrium is achieved when *β*_1_ < *β*_*c*_ < *β*_2_ and the state with *β*_1_ = *β*_2_ is unstable. *β*_*c*_ is the critical membrane excess area. *C*: two compartments of different base radii, where compartment 1 is smaller than compartment 2. The tension-area diagrams at constant intracellular pressure *P* are illustrated and *β*_*c*1_, *β*_*c*2_ are the critical *β* values for compartments 1 and 2 in *C*. As *Γ* scales with base radius, *a*, as *a*^2^, non-normalized tension *γ*_0_ is plotted in *C*.

If the two compartments have different base radii, the normalized pressure *p* scales as *a*^3^, while tension *Γ* scales as *a*^2^. For the smaller compartment, the critical normalized membrane excess area, *β*_*c*1_, is smaller than that for the larger compartment, *β*_*c*2_ (Fig. 4*C*). When the total membrane excess area *β*_1_+*β*_2_ is small, both compartments are on the left descending branches of their tension-area diagrams as in Fig. 5*A*. For a large total membrane excess area, *β*_1_+*β*_2_, the stable equilibrium is achieved when one compartment is on the left descending branch and another one is on the right ascending branch (Fig. 4*C*) of the tension-area diagram. The most stable state with the highest possible tension is achieved when the larger compartment is on its right branch (Fig. 4*C*).

**Fig. 5.**
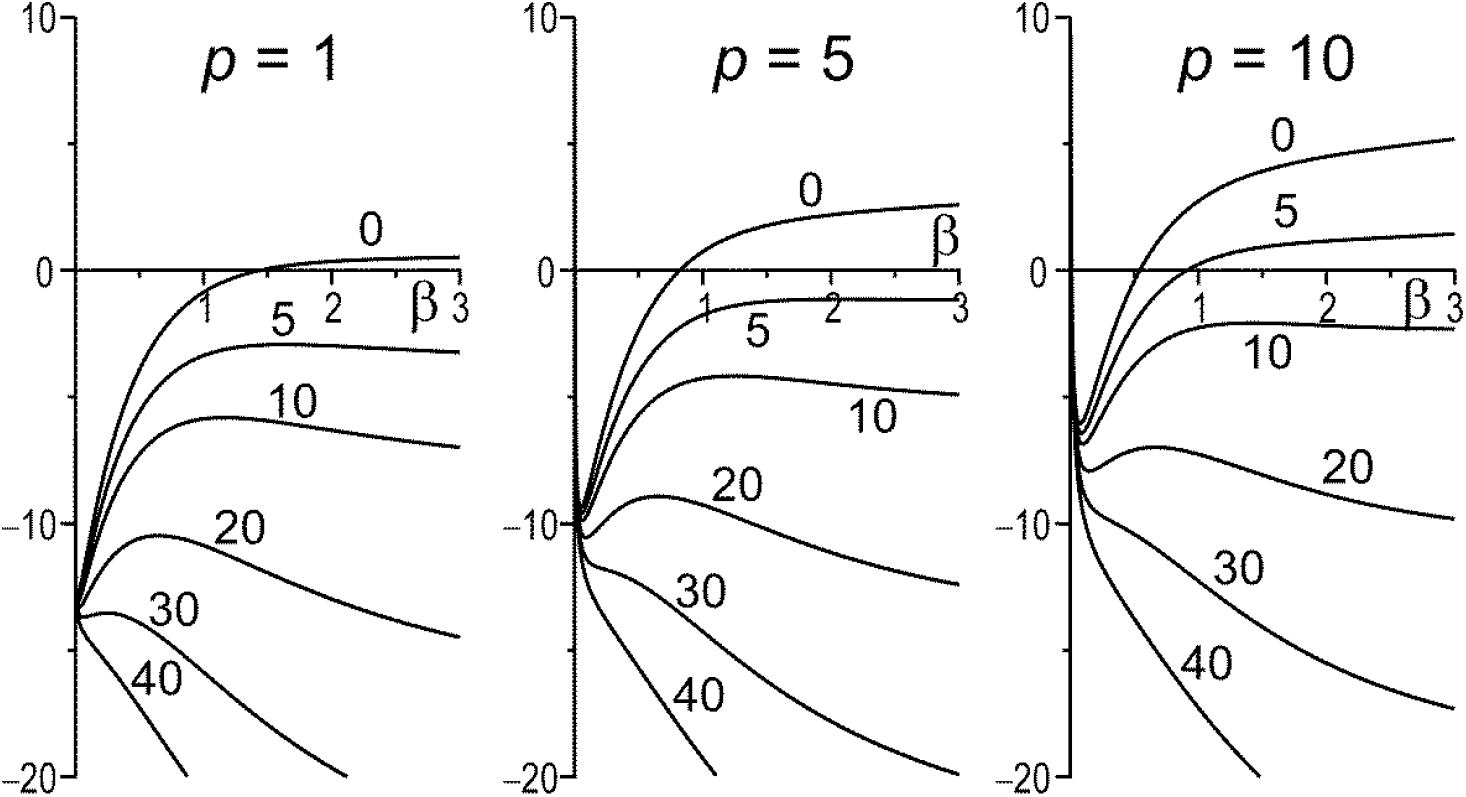
The dependence of the effective normalized membrane tension, *Γ*_*e*0_, on the normalized membrane excess area, *β*, at different normalized transmembrane pressure differences (shown on top of the plots) and the normalized protein concentration Θ*c*_0_ of 0, 5, 10, 20, 30, and 40 (shown next to the curves, respectively).

Now consider an ensemble of membrane compartments delineated with the picket fences of anchoring proteins between them. The total membrane area in the ensemble remains constant. Extracting membrane area from one compartment triggers membrane redistribution trough tension-driven flow across borders until a new steady state is established. In this state, membrane tension differs from its initial value, although it is the same in all compartments. The impact of membrane area reduction on steady-state membrane tension depends on the total membrane area.

If all compartments have equal base radii and the average membrane excess area is below its critical value, *β*_*c*_, the stable equilibrium corresponds to the descending branch of the diagram in all compartments, and the excess area is uniform. If the average excess area exceeds *β*_*c*_, the most stable equilibrium, state with the highest possible membrane tension, is achieved when all compartments except one are on the left descending branch of the tension-area diagram, and only one compartment accumulating almost all excess membrane area is on the right branch. For instance, at a moderate normalized transmembrane pressure of *p* = 1, in the steady state of an ensemble of 1000 compartments of identical size with an average membrane excess area of 0.1, one compartment would accumulate *β*>90, i.e. 90% of the total membrane excess area, while in all others, *β* < 0.01. If the compartments are of different sizes, the largest one accumulates most of the excess area of the entire ensemble.

### Osmotic pressure of membrane-dissolved proteins: a factor limiting membrane accumulation

As shown above, the mechanistic model predicts the “winner-takes-all” scenario in a compartmentalized cell membrane with sufficient excess surface area. The prediction appears somewhat paradoxical and lacks support from experimental observations suggesting that some unaccounted factors provide more uniform distribution of membrane excess area along cell surface.

A possible mechanism involves the presence of remnants of a sub-membrane elastic network within the compartment membrane, in addition to lipids and proteins. Notably, this membrane-reinforcing network should not be connected to the cortex at the compartment base but rather attached to the anchoring proteins and/or cortex at the compartment border. This reinforcement alters the tension-area diagram from a U-shaped to a Ͷ;-shaped curve, thereby limiting the accumulation of membrane area within a single compartment.

Another, and perhaps more plausible, mechanism that could resolve the “winner takes all” paradox is a difference in osmotic pressure among neighboring compartments. This difference is caused by variations in the concentration of membrane proteins in the compartments as lipid flow causing protein dissolution or concentration. When the protein concentration in neighbor compartments differs, the driving force for lipid flow between them is not solely the membrane tension but rather a combination of the mechanical tension and 2D osmotic tension.

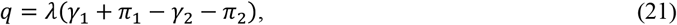

where π_1_, π_2_ are the 2D osmotic tensions of protein dissolved in the lipids in the compartments 1 and 2, respectively. Drawing an analogy between 3D osmotic pressure and 2D osmotic tension caused by membrane-dissolved proteins, express the osmotic tension as a function of protein concentrations in the compartments, *c*_*i*_, using van ‘t Hoff’s relation

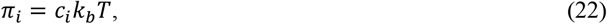

where *k*_*b*_ and *T* are the Boltzmann constant and absolute temperature; *c*_*i*_ is expressed in number of molecules per membrane area. In dimensionless form Eq. (22) gives

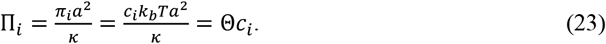

For *a* = 100 nm, κ = 10^19^ J, and *T* = 310 °K, obtain 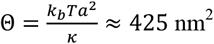.

For a typical protein concentration of 1 molecule per 50 lipid molecules (Quinn, Chapman 1991) and average membrane area per lipid molecule of 0.5 nm^2^ (Leftin et al. 2014), the obtained value for Π_*i*_ is approximately 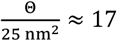. This value is substantial when compared to the characteristic normalised membrane tension caused by a realistic pressure difference at a reasonable membrane excess area, as depicted in Fig. 2.

The diffusion coefficient for lipids and lipid-dissolved proteins in cell plasma membrane were reported to 0.2÷1 μm^2^/s and (1.4÷3.6)·10^−3^ μm^2^/s, respectively (Kusumi et al. 2012). This means that the characteristic protein diffusion time, *t*_*D*_, at a distance of 2*a* ≈ 0.2 μm (i.e. the average width of a compartment), is 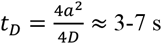. Assuming that the number of proteins in the *i*-th compartment, *n*_*i*_, is proportional to its base area, *A*_*bi*_: *n*_*i*_ = *c*_0_*A*_*bi*_, where *c*_0_ is a constant. Then, on a time scale ≤ *t*_*D*_, at which *n*_*i*_ remains nearly constant, obtain

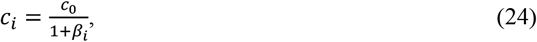

where *c*_*i*_ and *β*_*i*_ are the actual concentration of membrane-dissolved proteins and the membrane excess area in the *i*-th compartment.

The effective membrane tension in the *i*-th compartment that drives lipid area exchange with its neighbour compartments and included the osmotic component, *γ*_*e*_ is

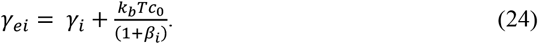

Membrane tension in a flat reservoir with a constant protein concentration, *c*_0_, that is equilibrium with the *i*-th compartment is

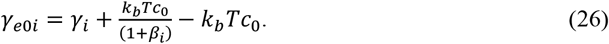

The *γ*_*e*0_ value is more convenient than *γ*_0_ for estimating membrane area redistribution between neighbor compartments at different concentrations of membrane-dissolved proteins, *c*_0_. The normalized *γ*_*e*0*i*_ value is

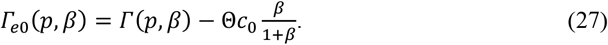

Within the ranges 0 < *β* < 1, 0 < *p* ≤ 30, and under stiff boundary condition, the numerically calculated tension-area diagram (Fig. 2*A*) – representing the dependence of Γ on *β* and *p* – can be approximated by a formula

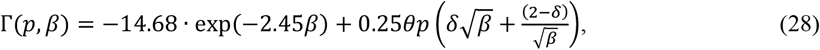

where *θ* = 0.63 and δ = 0.75 are fitting constants. The accuracy of the approximation is within ±1 for the specified *β* and *p* range. Notably, when *θ* = 1 and δ = 1, the second term of Eq. (28) matches the solution for torque-less membrane that obeys the Laplace formula of spherical dome (see Fig. 3*B*). The first term is an approximation of the calculated data at *p* = 0 whrere 14.68 is the normalized critical negative tension, 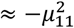,where μ_11_ is the 1-st root of the Bessel function *J*_1_ (Dharan et al., 2025).

The plots of *Γ*_*e*0_ for a compartment with stiff boundary conditions as a function of membrane excess area, *β*, at different normalized pressures *p* and values of a dimensionless parameter, Θ*c*_0_, calculated using Eq. (27) with the approximation Eq. (28), are shown in Fig. 5.

An increase in the osmolality factor Θ*c*_0_ results in a change in the effective tension area diagram from the U-shaped to Ͷ-shaped. Further increase in Θ*c*_0_ transforms the diagram into monotonic and L-shaped curve. These transitions occur at lower Θ*c*_0_ values at lower transmembrane pressure, as depicted in Fig. 5.

At any given normalized pressure difference, *p*, there is a range of the normalized protein concentration, Θ*c*_0_, for which the tension area diagram becomes Ͷ-shaped. At very low Θ*c*_0_, the diagram is U-shaped as was shown above in the absence of membrane-dissolved proteins. On the contrary, at very high normalized protein concentration, Θ*c*_0_, the diagram becomes L-shaped (Fig. 5). When the diagram is Ͷ-shaped, the normalized effective membrane tension, *Γ*_*e*_, has a local minimum at *β* = *β*_*c*_ and a local maximum at *β* = *β*_∗_. Respectively, at a given normalized transmembrane tension *p* there are effective tension values, *Γ*_*ec*_, *Γ*_*e*∗_, corresponding to *β*_*c*_ and *β*_∗_.

### Cortex-induced changes in membrane and cortex tension

Cortex deformations accompanied by changes in the compartment base area lead to a change in the normalized membrane excess area of a compartment and, therefore in membrane tension within this compartment. At a constant pressure difference and the total membrane area in a compartment, the dependence of membrane tension on the compartment base area is Λ-shaped if the tension – excess membrane area diagram is U-shaped and N-shaped if it is Ͷ-shaped. At a low membrane excess area, an expansion of a compartment base leads to an increase in the membrane tension and, vice versa, its shrinkage causes a decrease in the tension. At higher membrane excess area in a compartment that corresponds to the ascending part of the tension-area diagram, the base area and membrane tension change in opposite directions: base expansion leads to a decrease in tension while its shrinkage causes increase in tension.

### Membrane area distribution and tension in an ensemble of compartments with Ͷ-shaped tension-area diagram

The change from the U-shaped to Ͷ-shaped membrane tension-area diagram brings about a qualitative shift in the distribution of membrane surface area within an ensemble of interacting compartments. Membrane flow occurs from a compartment with lower effective membrane tension to its neighbor with higher effective tension. Assuming uniform compartment size and the protein density per base area, the effective tension-area diagram is Ͷ-shaped in all compartments as depicted in Fig. 6. In a steady-state equilibrium at the total excess membrane area below *β*_#_, the normalized excess area is the same in all compartments and the total effective tension-area diagram for the ensemble follows the steep descending branch of the diagram for a single compartment shown in blue in Fig. 6.

**Fig. 6.**
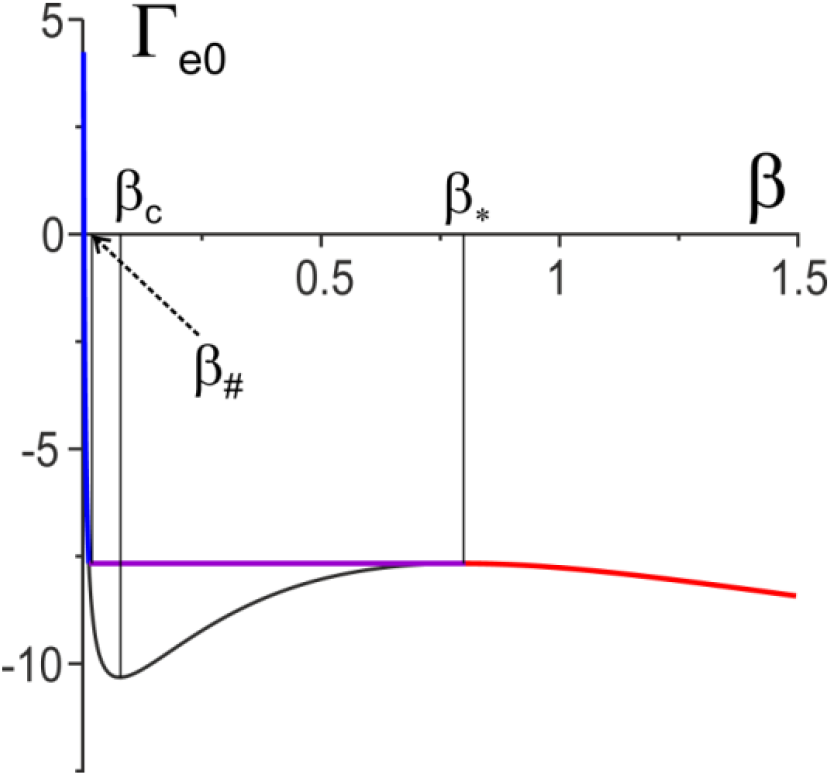
Effective normalized tension-area diagram for an ensemble of many Ͷ-shaped compartments at *p* = 5 and Θ*c*_0_ = 17. The critical, *β*_*c*_, peak, *β*_∗_, values of the normalized excess membrane area are shown as well as its switch value, *β*_#_, at which Γ_*e*_(*β*_#_) = Γ_*e*_(*β*_∗_) = *Γ*_*e*∗_. The steep descending branch, plateau, and a gentle descending branch are shown in blue, purple, and red respectively.

At a higher total membrane area, *β*_0_, certain compartment, termed “winners”, reside on the local maximum *Γ*_*e*_ = *Γ*_*e*∗_ of the diagram with *β* = *β*_∗_. Conversely, other compartments, labeled as “losers”, are positioned on the left descending branch of the diagram with low membrane excess area, *β*_#_, and the same constant effective tension, *Γ*_*e*∗_. The effective membrane tension is unaffected by a change in the total membrane excess area from *β*_#_ to *β*_∗_. The effective normalized membrane tension in all compartments remains constant and equal to *Γ*_*e*∗_ while the fractions of “winners” and “losers” change (purple line in Fig. 6). The effective tension-area diagram remains flat unless the total normalized membrane excess area exceeds *β*_∗_. Further increase in *β* results in a uniform distribution of the area across all compartments at a lower effective tension; as shown on the gently descending right red branch of the diagram in Fig. 6.

As a result, the tension-area diagram for the ensemble of a large number of Ͷ-shaped compartment of identical size has three distinct regions: a steep descending branch at *β*_0_ < *β*_#_, a plateau at *β*_∗_ < *β*_0_ < *β*_#_, and a less steep descending branch at high normalized membrane excess area *β*_0_>*β*_∗_.

If the Ͷ-shaped compartments of different sizes form an ensemble exchanging membrane area across the boundaries between them, the tension-area diagram for the ensemble slightly differs from the steep-flat-shadow diagram shown in Fig. 6. The flat middle part of the diagram becomes a descending line with a small although not zero inclination.

### Ensemble of compartments with Ͷ-shaped tension-area diagram and different base size

If the compartments in the ensemble have different sizes, the membrane redistribution becomes more complex. To assess the problem, we considered an ensemble of compartments with 15 different discrete radii of *a* = 30, 40, 50, …, 170 nm. The fractions of the compartments of different sizes followed a Gaussian distribution with a mean radius of *a*_*m*_ = 100 nm and a standard deviation of 35 nm, as estimated experimentally (Kusumi et al. 2012). Figure 7 illustrates the dependence of the normalized effective membrane tension on the effective normalized membrane excess area for these compartments at three different normalized pressures and at a constant normalized protein concentration of Θ*c*_0_ = 25.

**Fig. 7.**
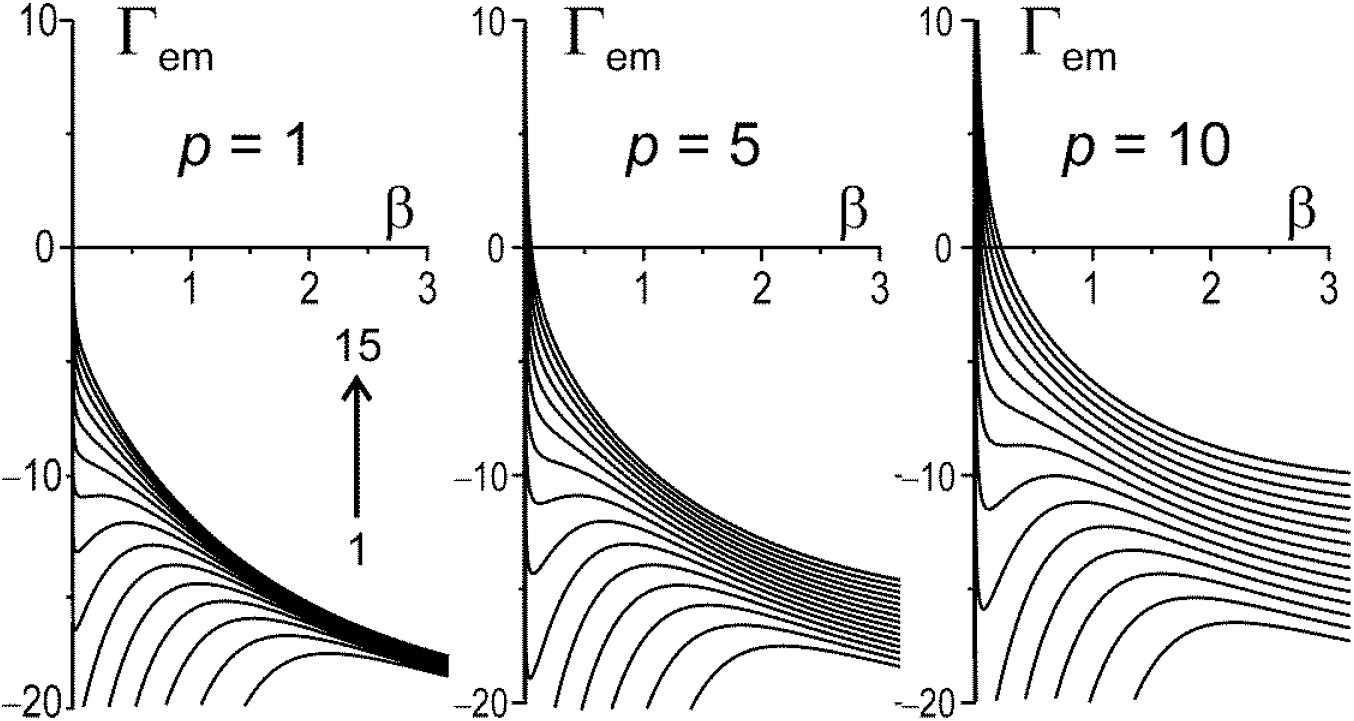
The effective normalized tension-area diagrams for membrane compartments of different size. The compartment radii *a* were 30, 40, 50, …, 150, 160, and 170 nm (from top to bottom, curves numbered 1, 2, …, 14, 15 respectively). Normalized pressure, *p*, on top of the plots and values of normalized effective membrane tension, *Γ*_*em*_, on the vertical axis correspond to a compartment of *a*_*m*_ = 100 nm. The normalized protein concentration, Θ*c*_0_, was 25 for all plots.

The plots shown in Fig. 7 were generated using Eq. (27)-(28) for various compartment radii. The shape of the effective tension-area diagram is influenced by both compartment size and the transmembrane hydrostatic pressure difference. For smaller compartments, the diagrams exhibit and Ͷ-shape, whereas for larger compartments, they display an L-shape. As the pressure difference increases, the transition from a Ͷ-shaped to L-shaped diagram occurs in smaller compartments (Fig. 7).

Next we modelled the distribution of membrane area between compartments of difference sizes, assuming that all compartments of the same size have the same normalized membrane excess area. The dependence of the effective tension in the all compartments that was established in the equilibrium steady-state on the total normalized membrane area is shown in Fig. 8.

**Fig. 8.**
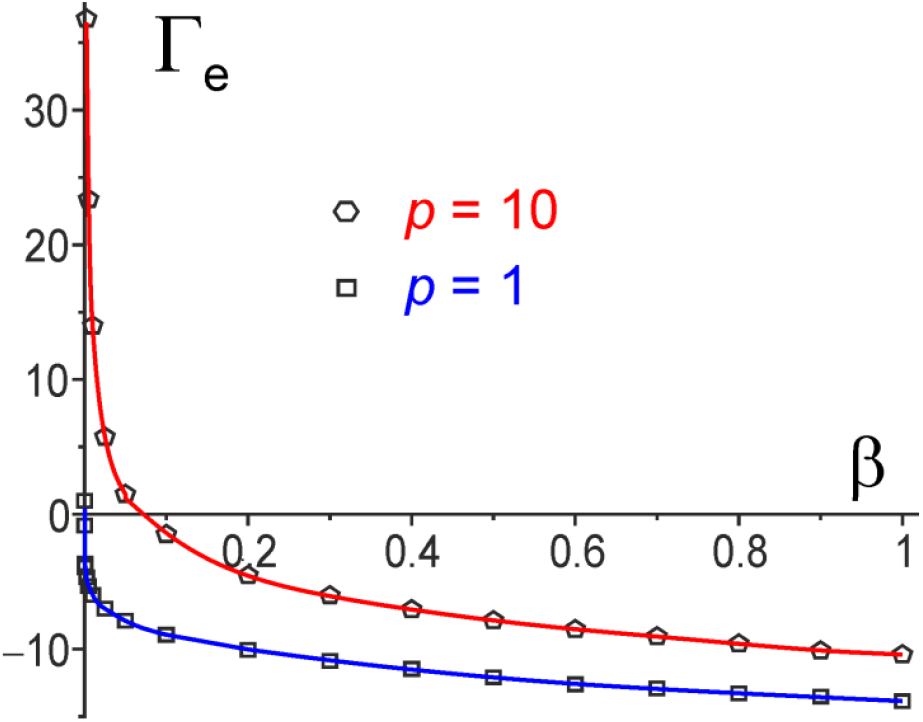
The effective normalized tension-area diagrams for an ensemble of compartments with radii ranging from 30 nm to 170 nm shown in Fig. 7 at two different normalized transmembrane hydrostatic pressures, *p*, 1 and 10. The *p* and Γ_*e*_ values on the plots correspond to compartment radius of *a*_*m*_ = 100 nm. The dimensional pressure (*P*) and effective tension (*γ*_*e*_) values were the same constant across all compartments. Respectively, the normalized values of *p* and Γ_*e*_ differ in compartments of various sizes.

In this ensemble of compartments, a higher normalized membrane excess area is evident in larger compartments compared to smaller ones, irrespective of the tested β0 values. However, it is important to emphasize that this accumulation is confined, primarily the total excess membrane area concentrated in compartments of medium to high radii.

The variability in compartment sizes leads to a smoothening of the characteristic “steep-flat-ramp” shape observed in effective tension-area diagrams for an ensemble of identical-sized compartments depicted in Fig. 7. The slope of these diagrams diminishes with an increase in the total normalized excess area. Notably, at higher pressure differences, the slope of the diagram is higher at any given *β*_0_ (Fig. 8).

## Discussion

### Pressure, tension and osmotic tension values

Before delving into potential biological implications of the model, it proves beneficial to estimate numerical values for membrane tension, *γ*, and pressure difference, *P*, in relation to their normalized counterparts. Assuming a compartment radius, *a*, of 100 nm, half the compartment size estimated by two different experimental methods (Kusimi et al. 2012), and a membrane bending stiffness, *κ*, of 10^−19^ J (Steinkühler et al. 2019), *Γ* values of 1. 10 and 50 correspond to γ values of 0.01, 0.1 and 0.5 mN/m, respectively which are within the range of the membrane tension values estimated experimentally (Gauthier et al. 2012; Lieber et al. 2013). Normalized characteristic osmotic pressure, Π, at protein-lipid molar ratio of 1:50 (Quinn, Chapman 1991) is about 17 that corresponds to a substantial osmotic pressure of π of ≈0.17 mN/m. Similarly, *p* values of 1 and 10 correspond to *P* values of 100 Pa and 1000 Pa, respectively. These values fall within the range of experimentally observed pressures (Chengappa et al. 2018).

### Comparison of model predictions with experimental observations

The model proposes a U-shaped mechanical tension-area diagram for a membrane compartment lucking remnants of cytoskeletal proteins and neglecting osmotic pressure caused by membrane-dissolved proteins (Fig. 2). At high pressure, when membrane bending stiffness becomes negligible, this behavior resembles that of a soup film being blown by pressure from a round aperture. When the pressure is zero, the flat membrane resists compressing tension until a threshold tension level is reached. Once the threshold is exceeded, the membrane buckles, and the post-buckled tension required to accommodate the excess membrane area in the compartment becomes lower than the threshold. Further increasing the membrane area within the compartment requires less compressive tension. These combined mechanisms produce the characteristic U-shaped tension-area diagram.

For an ensemble of many membrane compartments separated by picket fence borders, the membrane lipid flow through the fence between two compartments depends on both mechanical and osmotic pressure difference. The osmotic tension is generated by proteins dissolved within the lipid membrane within a compartment. If the borders are significantly less permeable to proteins than to lipids, the redistribution of membrane area between the compartments is constrained by protein concentration or dilution. This leads to a shift in the effective tension-area diagram from a U-shaped to a Ͷ-shaped (Fig. 5).

Another mechanism that may limit membrane area accumulation is the presence of cytoskeleton remnants bound to the compartment membrane and cortex at its border. Spectrin-associated proteins, such as ankyrin B, and protein 4.1 (nonerythroid isoform), have been found on the inner side of the membrane of expanding blebs (Charras et al., 2006). These proteins can elastically resist excessive membrane area accumulation and convert a U-shaped diagram into a Ͷ-shaped or an L-shaped one. Cortex reformation on the inner side of the membrane, which occurs within the first tens of seconds after bleb initiation (Charras et al., 2006), may similarly take place on the inner membrane of a compartment as its dome area expands due to membrane accumulation from neighboring compartments. The reformation of the cortex may further limit the membrane area in a compartment, preventing a “winner takes all” scenario.

In the steady-state equilibrium of an ensemble of many compartments with a U-shaped or Ͷ-shaped tension-area diagram and moderate or high total membrane excess area, the model predicts that larger compartments should accumulate more membrane excess area. Uneven roughness distribution, including domes as well as blebs and microvilli, was observed with scanning electron microscopy in different cells with large membrane excess areas (Erickson, Trinkaus 1976). A heterogeneous topology with submicron roughness of up to 100 nm in height was observed with scanning ion conductance microscopy (SICM) in live cells (Adler et al. 2010; Parmryd, Onfelt 2013; Gesper et al. 2020).

If the compartments with the Ͷ-shaped diagram in an ensemble are of similar size, the model predicts that at equilibrium the effective-tension-area diagram will have a characteristic “steep-flat-sloping” shape (Fig. 7). In other words, within a certain range of membrane area, the ensemble behaves as a membrane “reservoir” that can accumulate or release membrane at constant tension. This behavior was observed in experiments with membrane tethers pulled with an optical trap at a constant velocity: tension increased quickly when the pulling started, then remained constant as the reservoir was utilized, and then increased rapidly once the reservoir was depleted as in experiments of Raucher and Sheetz (1999).

### Membrane blebs as large compartments

Blebs are bubble-like protrusions of the cell membrane produced by intracellular pressure due to partial detachment of the cell membrane from the underlying cortex, cortex rupture, or active contraction (Charras, 2008). They play crucial roles in various cellular physiological and pathological processes including cell motility (Charras, 2008; Charras, Paluch 2008; Garcia-Arcos et al. 2024). During expansion, a bleb recruits membrane surface area from folds, bumps, and possibly structures such as caveolae (Parton et al. 2021) or microvilli (Morales et al. 2023). Bleb growth occurs rapidly for several seconds, then slows down, eventually ceasing after approximately 30 seconds, followed by retraction over 2–3 minutes (Charras et al. 2006; Charras 2008; Tinevez et al. 2009). Notably, bleb growth stops before cortex reassembly is fully completed, although some cortex-associated proteins appear on the inner surface within a few seconds after bleb begins to grow (Charras et al. 2006). Retraction is accompanied by the complete reassembly of the cortex, followed by its active contraction (Charras et al. 2006).

Under conditions that promote cell blebbing, multiple blebs with radii of several μm are often observed. Some of these blebs are very closely spaces and even can touch each other (Charras et al. 2006; Charras, Paluch 2008; Charras 2008). The total surface area of the multiple blebs can be comparable to the total cell cortex area (Charras, Paluch 2008; Charras 2008). The rapid growth of a bleb induced by local laser ablation of the cortex halts before it accumulates a substantial fraction of excess membrane area from the rest of the cell surface. This is evident from experiments with subsequent cortex ablation at a locus distant from the first bleb that results in the formation of another bleb of a similar or smaller size (Tinevez et al. 2009).

A growing bleb can be considered a membrane compartment with a border radius of approximately 0.5 μm which is about 5 times larger than that of a typical compartment. Therefore, as suggested by Eq. (12), at constant pressure difference, *P*, the reservoir membrane tension, *γ*_0_, and membrane bending stiffness, κ, the normalized pressure, *p*, and tension, Γ, of a bleb are ∼125 and ∼25 times higher, respectively, than those for a regular compartment of a radius 0.1 μm. The contributions of membrane bending stiffness to the shape of a bleb and membrane tension are negligible compared to those in a smaller regular compartment, except narrow region near the bleb border. The blebs are typically more than a hemispherical and have shape similar to that shown in Fig. 2C when the normalized membrane excess area is high.

Observations of blebbing cell show a tendency for preferential membrane area accumulation in blebs with larger border diameter, as predicted by the model developed above (Charras, Paluch 2008; Charras et al. 2006). However, the “winner takes all” scenario is not typically observed, indicating that some factors limit membrane area accumulation in a bleb. These factors may include remnants of cytoskeleton proteins in the bleb membrane connected to the underlying cortex on the bleb border or the osmotic pressure of membrane-dissolved proteins in the bleb membrane, as discussed above.

### Possible contribution of crumpled compartments to membrane reservoir

The tension dependence on the area of compartmentalized plasma membrane is likely dynamic, as membrane-dissolved proteins, although slowly, can still diffuse through the picket fences between compartments. The equilibration of protein concentration tends to convert the Ͷ-shaped tension-area diagram back to a U-shape. Conversely, the remnants of cytoskeletal proteins and cortex reformation on the inner surface of compartments with large membrane area limit membrane area accumulation, converting the U-shaped diagram to the Ͷ-shaped of L-shaped one. Our modeling suggests that an ensemble of membrane compartments of equal radius with a Ͷ-shaped tension-area diagram behaves as reservoir, maintaining nearly constant tension in a range of excess membrane area (Fig. 7). For a realistic distribution of compartment sizes, the resulting tension-area relationship of the ensemble is L-shaped with a long, gently descending branch (Fig. 8).

The excess area of plasma membrane is distributed among various structures, including protrusions formed by bundled actin filaments that push membrane outward (Morales et al. 2023), and membrane invaginations such as caveolae caused by membrane-bound proteins (Parton et al. 2020). These structures have a relatively high buffering capacity, capable of releasing a membrane area equivalent to tens of percent of the cell surface area (Sinha et al. 2010; Nassoy, Lamaze 2012). However, releasing this membrane area from these structures may require significant tension (Sinha et al. 2010; Nassoy, Lamaze 2012). Experiments involving repetitive tether pulls from cells showed an increase in the plateau length on the tension − tether length plot (Raucher, Sheetz 1999). These data demonstrated that high membrane tension, which causes a micro-bead to escape from the optical trap, increases the membrane area reservoir available for successive tether pulls. Our modelling suggests that crumples in the compartmentalized plasma membrane can form an essential part of the membrane reservoir. The flat or gently descending tension-area relationship for an ensemble of membrane compartments with Ͷ-shaped diagrams may also contribute to the slow tension propagation along as shown by Dharan et al. (2025).

### Model limitation

For simplicity, we considered only axisymmetric membrane compartments, although their shape can vary significantly (Kusumi et al. 2005, 2011, 2012). We believe that the simplification does not qualitatively affect the results, though it may introduce some quantitative differences.

However these differences are likely within the accuracy of the estimates of the model parameters values.

## Acknowledgements

The work was partially supported by the Israeli Ministry of Aliyah and Integration. The author extends deep gratitude to Professor Michael Kozlov for introduction to the biophysics and biomechanics of biological membranes as well as for invaluable discussions. Special thanks are due to Dr. Ben Zucker for his generous assistance in mastering the Surface Evolver software.

## Competing interests

The author declares no competing interests.

## Notes

### Competing Interest Statement

The authors have declared no competing interest.

